# Gene-to-Gene Coordinated Regulation of Transcription and Alternative splicing by 3D Chromatin Remodeling upon NF-κB activation

**DOI:** 10.1101/2023.08.07.552259

**Authors:** Paul Marie, Julien Ladet, Matéo Bazire, Lamya Ben Ameur, Sanjay Chahar, Nicolas Fontrodona, Tom Sexton, Didier Auboeuf, Cyril F. Bourgeois, Franck Mortreux

**Author notes:** [Lamya Ben Ameur], Vollum Institute, Oregon Health & Science University, Portland, OR 97239, USA.

## Abstract

The p65/RelA factor of NF-κB plays a pivotal role in coordinating gene expression in response to diverse stimuli, including viral infections. At the chromatin level, p65/RelA regulates gene transcription and alternative splicing (AS) through promoter enrichment and genomic exon occupancy, respectively. However, the mechanisms underlying the coordination of these processes across distinct genes remain elusive. In this study, we employed the HTLV-1 Tax oncoprotein, a potent activator of NF-κB, to investigate the integrative relationship between 3D chromatin architecture, NF-κB-regulated transcription and AS. Our analysis revealed that Tax induces a pronounced reorganization of the 3D genome, resulting in the formation of multigene complexes that comprise genes regulated either transcriptionally or through AS. Notably, we found that the Tax-induced gene-gene contact between the two master genes *NFKBIA* and *RELA* is associated with their differential regulation in gene expression and AS, respectively. Through dCas9-mediated approaches, we demonstrated that *NFKBIA*-*RELA* interaction is required for AS regulation and is caused by an intragenic enrichment of p65/RelA on *RELA*. Our findings shed light on new regulatory mechanisms upon HTLV-1 Tax and underscore the integral role of p65/RelA in coordinated regulation of NF-κB-responsive genes at both transcriptional and AS levels in the context of the 3D genome.

## INTRODUCTION

The process of gene expression involves multiple steps, including transcription and splicing, that work together to shape the transcriptome and ultimately determine cell behavior. The gene-to-gene coordination between transcription and alternative splicing events is thus required for proper cell response to stimulus. Cellular pathogens such as viruses can disturb this coordination, leading to both quantitative and qualitative alterations in transcriptome diversity, which can ultimately disturb cellular homeostasis and promote viral replication and persistence. A better understanding of mechanisms involved in coordinating transcription and splicing is thus necessary to comprehend the cellular response to viral infections.

The retrovirus Human T-cell Leukemia Virus type 1 (HTLV-1) primarily infects T lymphocytes, leading to a range of diseases, including adult T-cell leukemia/lymphoma (ATLL) and various inflammatory diseases (1). HTLV-1 replication initially involves cell-to-cell transmission followed by clonal expansion of infected cells which is responsible for the persistence of the virus and associated diseases (2). One of the key viral proteins involved in HTLV-1 infection is the oncoprotein Tax that has been first described as a transcriptional activator interacting with a variety of host cell proteins, including the NF-κB family of transcription factors. By hijacking NF-κB signaling, Tax deeply affects crucial mechanisms involved in immune response, cell survival, and inflammation, and plays a critical role in HTLV-1 pathogenesis (3).

At the molecular level, Tax interacts with multiple NF-κB and IκB proteins, including p65/RelA (4), p50 (5), c-Rel (4), p100 (6–8), p105 (9), IκBα (10,11), and NEMO (10,12). Tax activates the NF-κB pathway by promoting the degradation of the inhibitory protein IκBα, which controls the oscillating dynamics of NF-κB signaling (13,14). Tax induces a constitutive nuclear translocation of p65/RelA, resulting in chronic activation of target genes. Through post-translational modifications (15), Tax is able to form nuclear bodies enriched in NF-κB p65/RelA and splicing factor SFRS2/SC35 that have been proposed to participate in Tax-mediated activation of gene expression via NF-κB pathway (16,17). In this setting, we have recently demonstrated that when Tax is expressed, p65/RelA not only impacts promoters but is also enriched at gene bodies and induces alternative splicing changes, thereby broadening the influence of NF-κB on transcriptome diversity, both quantitatively and qualitatively (18). The transcription factor p65/RelA interacts with various splicing factors (18–22), including DDX17 which regulates splicing of GC-rich exons when recruited by p65/RelA in close proximity to the genomic target exon (18). It is noteworthy that Tax-induced alternative splicing primarily impacts genes that are not modified at the gene expression level (18,23). Further, experimental tethering of p65/RelA to its genomic target exon is sufficient to locally affect alternative splicing, indicating that p65/RelA acts on splicing regardless of its activity on the target gene promoter (18). Given the tightly controlled coordination of NF-κB-regulated genes to mediate cellular functions and responses to stimuli, these findings raise questions whether these two distinct activities of p65/RelA are coordinated.

The transcriptional coordination of NF-κB-responsive genes is regulated through a complex interplay of mechanisms that includes the oscillating dynamics of NF-κB signaling (13,14,24), the differential DNA affinity of homo- or heterodimers formed by NF-κB subunits (25,26), the recognition of κB sites and flanking sequences (26–28), as well as specific co-factors and post-translational modifications of NF-κB (29,30). More recently, the 3D conformation of chromatin has emerged as a critical determinant of NF-κB-mediated gene expression regulation (31). NF-κB p65/RelA is able to promote DNA looping through interactions with various transcription and chromatin remodeling factors (32,33). NF-κB signaling upon TNF-α has been associated with the formation of specialized transcription factories, also called NF-κB factories, where responsive-coding and -miRNA genes are transcribed in a coordinated manner (34). Such long-range interactions of co-transcribed genes can occur in a hierarchical relationship that contributes to the temporal coordinated regulation of responsive genes (35).

Of note, 3D chromatin conformation has also been involved in co-transcriptional alternative splicing regulation, in combination with other various factors such as RNA binding proteins, elongation rate of transcription, and chromatin remodeling factors (33–41). Active chromatin has been observed highly concentrated in the nuclear center and around nuclear speckles (NS), which are dynamic subnuclear structures enriched in splicing factors and other RNA processing proteins (42). NS are enriched in GC-rich genomic sequences in their close vicinity (43). In particular, it appears that exon-intron units with high-GC content are preferentially localized in the nuclear center in association with NS, compared to their low-GC counterparts (44,45). Importantly, high- and low-GC exon-intron units display different splicing patterns, and their experimental shifting from one genomic location to another affects both their transcription and splicing regulation, indicating that the nuclear location of the gene determines its regulation in alternative splicing (45). In line with this, the formation of multigene complexes via trans-chromosomal contacts in foci enriched in splicing factor RBM20 has been shown to function as a splicing factory, suggesting that nuclear condensates may act as hubs regulating both transcription and alternative splicing of cell-signaling responsive genes (46). The question thus arises whether 3D chromatin conformation could play a role in coordinating transcription and alternative splicing events promoted by p65/RELA upon Tax expression.

Here, we provide molecular evidence that Tax induces drastic changes in the 3D conformation of the cell genome that are accompanied by differential gene expression and alternative splicing modifications. In particular, we show that a trans-chromosomal interaction involving *NFKBIA* and *RELA* genes is required for Tax-induced skipping of exon E6 of *RELA*, leading to the expression of an isoform of p65/RelA with a unique gene expression activation pattern. Engineering-based approaches via dCas9 demonstrate that *NFKBIA*-*RELA* contacts are caused by the DNA binding of p65/RelA at the close vicinity of the genomic *RELA* E6, without affecting target gene expression. These results provide insight into the interplay between HTLV-1, NF-κB signaling, and 3D genome conformation in reprogramming cellular function, demonstrating how the formation of a p65/RelA dependent multigene complex coordinates NF-κB-related transcription and alternative splicing events.

## MATERIAL & METHODS

### Cell culture and transfections

The human embryonic kidney (HEK) 293T-LTR-GFP cell contains an integrated GFP reporter gene under the control of the viral HTLV-1 long terminal repeat (LTR). HEK293T-LTR-GFP were cultured in DMEM + 4.5 g/L D-Glucose + Pyruvate + GlutaMAX™, supplemented with 10 % heat-inactivated FBS and 1 % Penicillin/Streptomycin. The transfection efficiency of cells with Tax, Tax-M22, and dCas9-T2a-GFP constructs was assessed by measuring GFP fluorescence with EVOS® FL Imaging System. RNA guides for dCas9 (sgRNA) (Supplementary Table 1) were designed using UCSC ‘CRISPR targets’ tracks and sequences with overall highest prediction scores and lowest off-targets. The sgRNAs were cloned into the BsmBI sites of the CRIZI plasmid (generously gifted by Philippe Mangeot, CIRI, Lyon, France). For dCas9-p65 cloning, WT-p65 was amplified from HEK-293T-LTR-GFP cDNA and cloned into the BamHI and XbaI sites of dCas9-empty-GFP plasmid (generously gifted by Reini F. Luco, Institut Curie, Orsay, France). The expression vectors pRELA-WT and pRELAΔE6 were produced using sequences amplified from HEK293T cDNA and the V5 sequence was added in 5’. Sequences were then cloned in pXJ41 plasmid using NotI and BamHI restriction sites. All sequences were verified by Sanger sequencing.

Transfections were performed using pSG5M-Tax-WT, pSG5M-Tax-M22, pLV hU6-sgRNA hUbC-dCas9-KRAB-T2a-GFP (addgene plasmid #71237), sgRNA-CRIZI, pBabe-Puro-IKBalpha-wt (Addgene plasmid #15290), dCas9-p65, dCas9-empty using Jeprime (Polyplus Transfection) following manufacturer’s instructions, and cells were harvested 48h post transfection.

HTLV-1-chronically infected lymphocytes C91PL and non-infected Jurkat (both kindly gifted by Renaud Mahieux CIRI, Lyon, France) were cultivated in RPMI 1640 medium (Gibco, Life Technologies) supplemented with 25 mM HEPES, 10 % heat-inactivated FBS and 1 % Penicillin/Streptomycin.

### RNA-seq analysis

RNA-seq data have been previously published (18) and deposited on NCBI GEO under the accession number GSE123752. Briefly, poly-A transcripts were extracted at 48h post-transfection from 293T-LTR-GFP cells transfected with pSG5M-Tax or pSG5M empty vectors. RNA-seq libraries were prepared from biological triplicates and sequenced using the illumina HiSeq 2500. Paired-end pairs of reads were then analyzed using the pipeline FaRLine and the FasterDB database (47). HTSeq-count (v0.7.2)(48) and DESeq2 (v1.10.1) (49) were used calculating gene expression level and for computing differential gene expression levels, respectively. The significance thresholds were set to 5% for ΔPSI (differential percentage of spliced-in sequence) (p < 0.01, Fisher’s exact test) and 0.6 for log2-gene expression changes (p<0.05, Wald test using Benjamini and Hochberg method), respectively. Relative genomic distance analysis between alternative splicing events (ASE) and differential gene expression was carried out using reldist from Bedtools. Control set of ASE was generated using the shuffle function of Bedtools.

### Hi-C assays and data processing

The Hi-C experiment and library preparation were conducted following the in situ method described by Rao and colleagues with slight modifications (50). Briefly, HindIII enzyme was used to digest cells overnight at 37°C with 500U, followed by biotin filling and proximity ligation for 4 hours at 18°C with 2000U T4 DNA Ligase. After reverse-crosslinking, DNA was purified and sheared to ∼300 bp fragments using Covaris S220 sonicator. The biotin-containing fragments were then immobilized on MyOne Streptavidin T1 beads. The NEXTflex adaptors were ligated, and the fragments were PCR amplified for 8 cycles using the KAPA HiFi Library Amplification Kit. Double-size selection was performed using AMPure XP beads to isolate fragments between 300 and 800bp. Sequencing of Hi-C libraries was performed on Illumina HiSeq4000 with 2×50 bp paired-end reads (Genomeast, IGBMC, Illkirch, France). The raw reads were independently mapped to the hg19 reference genome using the HiC-Pro pipeline (version v3.1.0), and the output statistics, including the total number of sequenced reads and the number of valid interactions after filtering, are presented in Sup Fig 1.

**Figure 1.**
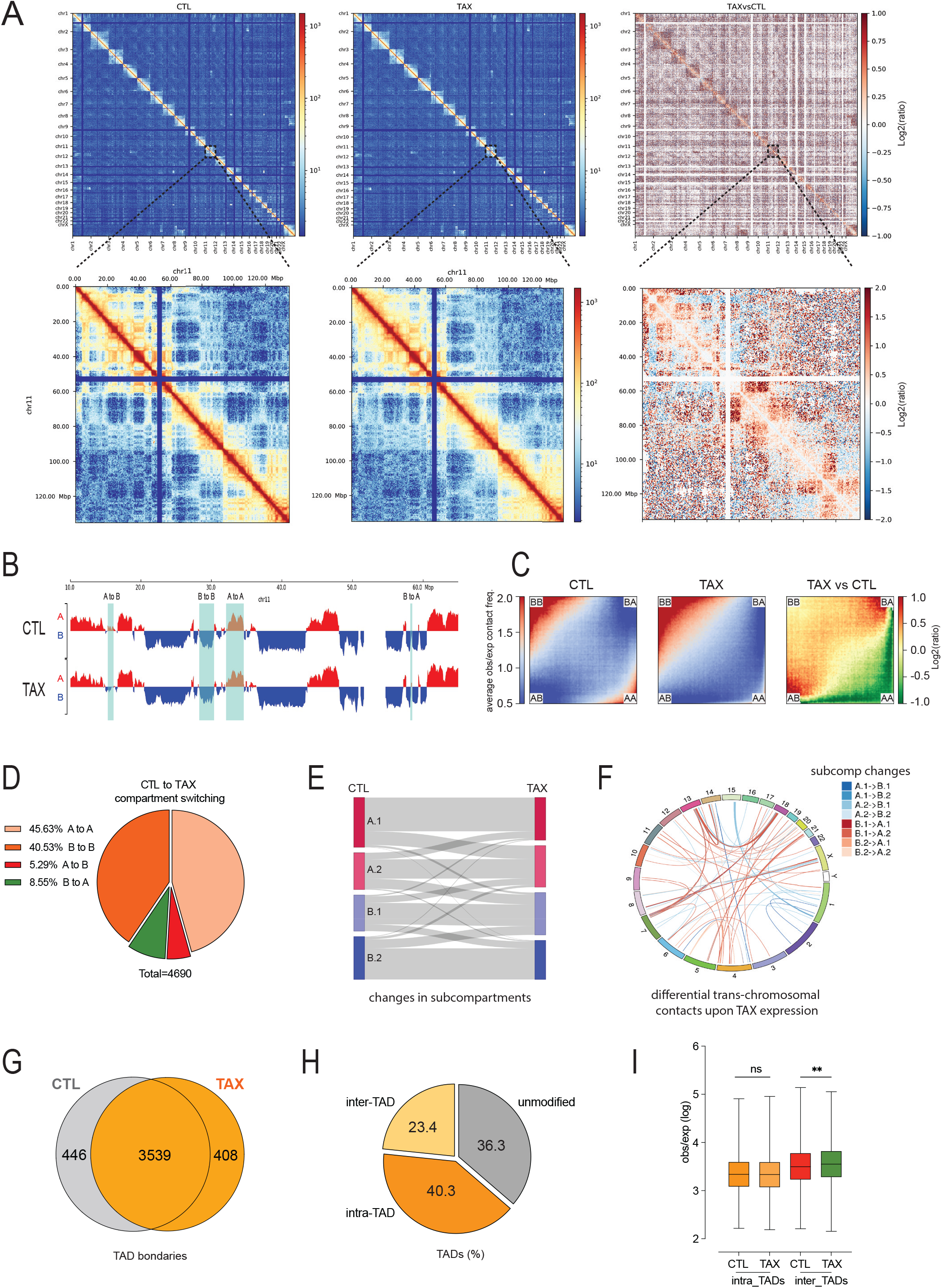
**Tax reshapes the 3D chromatin conformation of the genome.** **(A)** Heatmap showing the normalized Hi-C interaction frequency map of chromosomes in control cells (CTL) and cells expressing the Tax protein (Tax). The upper panel depicts all chromosome matrices with diagonal line representing the intra-chromosomal contacts, while the off-diagonal regions show the inter-chromosomal interactions. The lower panel zooms at 500 kb resolution across entire chromosome 11. The right panel represents the differential heatmap between Tax and CTL conditions, with the log2 ratio indicated by the color scale. **(B)** compartment A and B in both Tax and CTL conditions. Compartment A and B are respectively shown in red and blue. The panel demonstrates all possible switch between compartment A and B (cyan square). The x-axis shows the genomic coordinates, and the y-axis represents the normalized compartment score. **(C)** Saddle plot of Hi-C data showing the homotypic (A-A and B-B) and heterotypic (A-B and B-A) cis-interaction frequencies. The right panel shows the differential saddle plot between Tax and CTL in log2 ratio contacts. **(D)** Percentage of switching compartments in cells expressing Tax (Tax) compared to control cells (CTL). **(E)** Sankey plot showing the proportion of genomic segments that switch between each subcompartment in Tax and CTL conditions. **(F)** Circos plot showing differential trans-chromosomal contacts (log2FC≥2) identified by Hi-C data involved in the switching of subcompartments from control cells to Tax-expressing cells. **(G)** Venn diagram showing the overlap of TADs boundaries identified in control cells and cells expressing Tax. **(H)** Percentage of TADs showing differential intra-TAD and inter-TAD contacts upon Tax expression. **(I)** Boxplots of the observed over expected (obs/exp) interaction frequencies for intra-TAD and inter-TAD interactions in control cells and Tax expressing cells. The obs/exp ratios are expressed as logarithmic value.

The Hic explorer pipeline (v3.7.2) was used to process the Hi-C data and plot the matrices (51). The matrices were first normalized to each other (hicNormalize), then corrected (hicCorrectMatrix) for removing bins with low numbers of reads and balanced using ICE (52) at resolutions of 10k, 50k, 100k, and 500k bp. Corrected matrices were plotted (hicPlotMatrix) on a log scale at a resolution of 500k. The differential matrix was obtained using a log2ratio comparison between Tax and CTL cells on corrected matrices at the 500k resolution (hicCompareMatrices). The ratio between short-range and long-range contacts was computed using hicPlotSVL with the threshold distance set to 1e6 nucleotides.

The inference of A/B compartments and changes in compartmentation between Tax and CTL matrices were analyzed using the pipeline dcHiC with matrices at a resolution of 100k (53). The full genome compartment strength was calculated as (AA + BB) / (AB + BA) from the log2 ratio of observed versus expected contacts sorted by their eigen vector values (hicCompartmentalization). A saddle plot was generated with cooltools (https://github.com/open2c/cooltools) using cis-contacts from matrices at a resolution of 50k. The eight subcompartments were assigned by the pipeline

CALDER with matrices at a resolution of 10k and the binning variable set to 50k (54). The Deeptools’ PlotEnrichment (55) function was then used to validate CALDER predictions regarding their subcompartment enrichment levels in histone marks H3K27ac, H3K4me3, and H3K9me3 publicly available for HEK293T cells in ENCODE (ENCFF706GDY, ENCFF219COP, ENCFF800LFN, ENCFF063WXN, ENCFF626VTP, and ENCFF295FIO). Comparison of both compartment and subcompartment orientations between Tax and CTL conditions was carried out using Bedtools (56). Sankey diagram was built in R using the networkD3 package. Genomic regions corresponding to both subcompartment transition and inter-chromosomal changes upon Tax were obtained using bedtools intersect function with coordinates of Tax vs CTL log2ratio contact matrix computed at 50kbp and those of the subcompartment changes grouped as follows: A.1 (A.1.1 + A.1.2), A.2 (A.1.1 + A.1.2), B.1 (B.1.1 + B.1.2), and B.2 (B.2.1 + B.2.1). Gene ontology analysis and motif enrichment in promoters involved in 3D chromatin contacts in Tax expressing cells was performed with ShinyGO 0.76.3.

The identification of topologically associating domains (TADs) was carried out using the HicFindTADs function of the hicexplorer software with a matrix resolution of 50k. A minimum and maximum window length of 150k and 500k, respectively, were set for the analysis, and a q-value threshold of false discovery rate (FDR) was set to 0.05. The putative boundary and surrounding bins were differentiated by setting a minimum differential threshold (delta) to 0.01. The similarity of TAD boundaries between the control (CTL) and Tax-expressing cells was assessed by using the Bedtools’ Jaccard statistic with a tolerance window of 50 kbp. Intra-TAD and inter-TAD interactions were analyzed by merging the TAD domains obtained from both CTL and Tax conditions using the hiCMergeDomains function of Hicexplorer. The CTCF binding sites (ENCODE ENCFF895AOX) were used to delineate TAD boundaries, considering that boundaries within 20kbp of each other were merged as one. The merged TAD boundaries were then utilized to calculate the intra-TAD and inter-TAD differential contacts between Tax and CTL matrices (hicDifferentialTAD). Significant intra-TAD and inter-TAD interactions were defined thanks to the Wilcoxon rank-sum test with a p-value threshold set to 0.05. Bedtools intersect function allowed assigning gene deregulations to differential chromatin interactions between Tax vs CTL (log2(obs/exp) at 50 k). Network analysis of ASE, DGE, and chromatin contacts was performed with Cytoscape (3.9.1). Pile-up analysis was carried out with coolpup.py (57) using expected/observed signal (bin size 30kb) surrounding (1Mbp) the TAX-regulated genes. Randomly picked genes were obtained by bedtools (sample). The level of contact enrichment was estimated by the average value of interactions in the central 100 x 100 pixel square.

### Western Blot

Cells were washed twice with 1 X PBS, harvested and incubated with lysis buffer (10 mM Tris HCl pH 8, 50 mM NaCl, 5 mM EDTA, 1 % NP-40, 1 % SDS, 1 mM DTT, supplemented with protease inhibitors and 10 mM NaF) for 30 min on ice. Total proteins were sonicated on a Bioruptor Plus (Diagenode) (position high, 5 x 30’/30’ cycles), and cell debris were pelleted and discarded. 20 µg of total proteins were used for SDS-PAGE electrophoresis on NuPAGE™ 4–12% Bis-Tris Protein Gels and transferred on nitrocellulose membranes using the Trans-Blot® TurboTM Blotting System. Membranes were saturated in 1 X TBS-Tween containing 5% non-fat milk for 1 h and were incubated overnight at 4°C with primary antibodies diluted in 1 X TBS-Tween containing 5% non-fat milk. The primary antibodies used were the following: *RELA* (sc-109, Santa Cruz), Tax (1A3, Covalab), Vinculin (ab129002, abcam), IkBa (L35A5 Cell Signaling Technology). Membranes were washed three times in 1X TBS-Tween and incubated with HRP-conjugated secondary antibodies for 1 h before washing for three times. HRP substrate (Immobilon Forte (Millipore)) was deposed onto the membrane before reading chemiluminescence on a Chemidoc (BioRad).

### RNA extraction, PCR, and real-time quantitative PCR

Total RNAs were extracted using TRIzol (Invitrogen). 2.5 µg of total RNA was treated with dsDNase (Thermo Scientific) and retro-transcribed using Maxima First Strand cDNA Synthesis (Thermo Scientific) according to manufacturer’s instructions. PCRs were performed with GoTaq polymerase (Promega, Madison, WI, USA) using 7.5 ng cDNA as template, were run on agarose gel electrophoresis and read using Geldoc (BioRad). Quantitative PCR were performed with SYBR® Premix Ex Taq TM II (Tli RNaseH Plus) on LightCycler 480 II using 7.5 ng of cDNAs. Melting curves were checked for non-specific product absence. Relative expression levels were calculated using the ΔΔCt method on technical duplicates or triplicates and normalized to GAPDH expression. Alternative exons inclusion rates were calculated using ΔΔCt method and were normalized to a constitutive exon. Oligonucleotide sequences used are listed in Supplementary table 1.

### Chromatin immunoprecipitation

2×10^7^ cells were crosslinked with 1% formaldehyde for 10 min at room temperature. Crosslinking was quenched by addition of glycine to a final concentration of 0.125 M. Cells were harvested and incubated 30 min at 4°C in lysis buffer (1% Triton X-100, 0.1 % Sodium Deoxycholate, 0.1 % SDS, 140 mM NaCl, 50 mM HEPES-KOH pH 7.5, 1 mM EDTA, supplemented with protease inhibitor and phosphatase inhibitor). Nuclei were pelleted and resuspended in shearing buffer (10 mM Tris-HCl pH 8.0, 1 mM EDTA, 2 mM EDTA, 0.1% SDS, supplemented with protease inhibitor and phosphatase inhibitor). Chromatin was sheared to an average fragments size of 100-500 bp using Covaris S220 (Peak Power: 140; Duty Factor: 5; Cycles/burst: 200, 20 min). Immunoprecipitations were performed using 25 µg of chromatin diluted in dilution buffer (1% Triton X-100, 0.01% SDS, 150 mM NaCl, 10 mM Tris HCl pH 8, 1 mM EDTA, supplemented with protease inhibitor and phosphatase inhibitor), and incubated with 5 μg of anti-*RELA* (#36369, Active Motif), anti HA (H3663, Sigma-Aldrich) or control Rabbit or Mouse IgG (10500C and 10400C respectively, ThermoFisher) overnight at 4 °C under rotation. chromatin-antibodies complexes were recovered by incubation with 30 μl of Dynabeads® Protein A/G 1:1 mix (ThermoFisher) for 2 h at 4°C under rotation. Complexes were washed for 5 min at 4°C under rotation with the following buffers : Wash 1 (1% Triton X-100, 0.1% NaDOC, 150 mM NaCl, 10 mM Tris-HCl pH 8), Wash 2 (1% NP-40, 1% NaDOC, 150 mM KCl, 10 mM Tris-HCl pH 8), Wash 3 (0.5% Triton X-100, 0.1% NaDOC, 500 mM NaCl, 10 mM Tris-HCl pH 8), Wash 4 (0.5% NP-40, 0.5% NaDOC, 250 mM LiCl, 20 mM Tris-HCl pH 8, 1 mM EDTA), Wash 5 (0.1% NP-40, 150 mM NaCl, 20 mM Tris-HCl pH 8, 1 mM EDTA) and twice with Tris-EDTA buffer (10 mM Tris pH 8.0, 1 mM EDTA). The immunoprecipitated chromatin was eluted in elution buffer (1 % SDS, 200 mM NaCl, 100 mM NaHCO3) and crosslink was reversed by adding 100 µg proteinase K and incubating overnight at 65°C. Immunoprecipitated chromatin was purified by phenol-chloroform extraction, and quantitative PCR was performed using SYBR® Premix Ex Taq (Tli RNaseH Plus) (Takara) on a LightCycler 480 II (Roche, Mannheim, Germany), using manufacturer’s recommended thermocycling conditions. Values were expressed as fold enrichment relative to the signal obtained for the control IgG immunoprecipitation. Primers used for quantitative ChIP experiments are available in Supplementary Table 1.

### Chromosome Conformation Capture (3C)

1 x 10^7^ cells were cross-linked with 1 % formaldehyde (ThermoFisher) for 10 min at RT. Formaldehyde cross-linking was stopped by addition of glycine for 5 min to a final concentration of 0.125 M. Cells were washed twice with cold PBS and harvested. Cells pellets were lysed for 20 min on ice using lysis buffer (50 mM Tris HCl pH 7.4, 150 mM NaCl, 0.5 % NP-40, 1 % Triton X-100, 5 mM EDTA and supplemented with protease inhibitor cocktail). Nuclei were harvested by centrifugation and washed in 1.2 X restriction buffer (depending on the restriction enzyme used) and resuspended in 500 µl of the same buffer. For nuclear membrane permeabilization, SDS was added to a final concentration of 0.3 % followed by incubation at 37°C for 1 h under agitation (800 rpm). SDS was neutralized by addition of Triton X-100 to a final concentration of 2.5 % at 37 °C for 1h h at 900 rpm. Digestion was realized by addition of 50 U DpnII or ApoI at 37 °C for 3h at 900 rpm followed by another 50 U and overnight incubation at 37 °C at 900 rpm. Digestion efficiency was assessed on a reverse cross-linked and purified aliquot by agarose gel electrophoresis (data not shown). Restriction enzyme was inactivated by incubation at 65 °C for 20 min at 900 rpm. Digested chromatin was proximity-ligated by diluting restriction reaction in 7 ml of 1X ligation buffer. Ligation was performed by addition of 50 U T4 DNA ligase and 3h incubation at 16 °C at 750 rpm followed by another 50 U and overnight incubation at 16 °C at 750 rpm. Ligation efficiency was assessed on a reverse cross-linked and purified aliquot by agarose gel electrophoresis (data not shown). Cross-link was reversed by addition of 30 µg proteinase K and overnight incubation at 65 °C at 900 rpm. 3C templates were purified by phenol/chloroform extraction and ethanol precipitation.

3C-PCR were performed using semi nested primers (Supplementary Table 1) and two rounds of PCR. for the first round, 50 ng of 3C template was used and amplified using Phusion Green HSII High-Fidelity PCR Master Mix and a final primer concentration of 0.5 µM using the following program: 3 min of denaturation at 98°C, followed by 20× (98°C for 10 s, primer-specific annealing temperature for 30 s, 72 °C for 45 s) and 10 min final extension at 72 °C. The first-round of 3C-PCR was analyzed by gel agarose electrophoresis to check the absence of any significant amplification band before performing second-round 3C-PCR. Second-round 3C-PCR was performed using the same primer as the first round for *RELA* viewpoint, and a different primer closer to the restriction site for *NFKBIA* viewpoint. One µl of the first round PCR was amplified using Phusion Green HSII High-Fidelity PCR Master Mix and a final primer concentration of 0.5 µM using the following program: 3 min of denaturation at 98°C, followed by 25× (98°C for 10 s, primer-specific annealing temperature for 30 s, 72 °C for 45 s) and 10 min final extension at 72 °C. Optimal annealing temperatures were determined using Thermofisher Phusion tm calculator (www.thermofisher.com/tmcalculator). 3C-PCR products were analyzed by gel agarose electrophoresis.

### Circular Chromosome Conformation Capture (4C-seq) and data processing

For Circular Chromosome Conformation Capture, the same protocol as for 3C was performed. 50 µg of 3C template was digested by 50 U second restriction enzyme (Csp6I, ThermoFisher) in 1X Buffer B (ThermoFisher) for 2 h at 37 °C at 500 rpm. Digestion efficiency was assessed on a purified aliquot by agarose gel electrophoresis (data not shown). Restriction enzyme was inactivated by incubation at 65

°C for 20 min at 900 rpm. The second ligation was performed under diluted conditions by addition of 50 U T4 DNA ligase and 2 h incubation at 16 °C at 750 rpm followed by another 50 U and 2 h incubation at 16 °C at 750 rpm. Ligation efficiency was assessed on a reverse cross-linked and purified aliquot by agarose gel electrophoresis (data not shown). 4C templates were purified by phenol/chloroform extraction and ethanol precipitation followed by an additional JetSeq™ Clean purification.

4C sequencing primers (Supplementary Table 1) were designed as previously described (58) and tested on a small portion of 4C template with increasing templates amounts to determine linear range, efficiency, minimal primer dimer amplification, and optimal annealing temperature of each primer pairs. 4C libraries were prepared in two steps of PCR. The first step consisted in an amplification of ligation fragments specific of each viewpoint using 100 to 400 ng template depending on primer linearity. Expand™ Long Template PCR System (Roche) was used with the following thermocycling program: 2 min of denaturation at 94°C, followed by 16× (94°C for 15 s, primer-specific annealing temperature for 1 min, 68°C for 3 min) and 7 min final extension at 68 °C. All first step PCR primers are listed in Supplementary Table 1. Multiple PCR reactions were performed, pooled together and purified using JetSeq™ Clean purification according to manufacturer’s instructions. The second PCR step (used to add illumina adapters, indexes and sequencing sequences) was performed on 5 µl of purified first step PCR products using Expand™ Long Template PCR System (Roche) with the following thermocycling program: 2 min of denaturation at 94°C, followed by 20× (94°C for 15 s, 60 °C for 1 min, 68°C for 3 min) and 5 min final extension at 68 °C. All second step PCR primers are listed in Supplementary Table 1. 4C templates were purified using JetSeq™ Clean purification according to manufacturer’s instructions. Libraries quantity and purity were determined on a 2200 Tapestation (Agilent) with a D500 ScreenTape (Agilent). Sequencing of 4C libraries was performed on Illumina Nextseq 550 with 150 bp single-end reads (Genomeast, IGBMC, Illkirch, France).

Computational analysis of 4C-seq data was performed for each replicate and viewpoint using the fourSig pipeline (59) with the following parameters: window.size=5, iterations=1000, fdr=0.001, and fdr.prob=0.01. For each viewpoint, triplicate outputs were normalized to each other using multiBamsummary and Bamcompare from deepTools (55). Differential chromatin contacts between CTL and Tax conditions were determined using the DESeq2 R package (FDR=0.05). Figures were generated using pygenometrack (version 3.6) (60).

## RESULTS

### Tax reshapes the 3D genome

To test whether 3D chromatin conformation could play a role in coordinating transcription and alternative splicing events promoted by p65/RELA upon the viral oncogene HTLV-1 Tax, we first evaluated the impact of Tax on 3D chromatin organization. To this end, we performed an unbiased chromatin interaction mapping using Hi-C analysis of HEK293T cells with and without Tax expression. Hi-C data were analyzed using the HiC-pro pipeline (61), which ensured high-quality experiments as evidenced by high mapping rates (Sup Fig 1). We identified over 160 million interactions, including over 20 million trans-chromosomal contacts, in both control (CTL) and Tax-expressing cells (Sup Table 2). Consistent with previous studies on chromosome territories (62,63), the Hi-C contact maps demonstrated higher frequencies of cis-versus trans-chromosomal interactions (Figure 1A). While the Hi-C maps from CTL and Tax-expressing cells showed similar bulk properties (Figure 1A and Sup Fig 2), we noticed a decrease in short-range (<1 Mb) and an increase in long-range (>1 Mb) interaction frequencies upon Tax expression (Sup Fig 3 A-B). These changes were particularly evident in differential heatmaps (Figure 1A, right panel).

**Figure 2.**
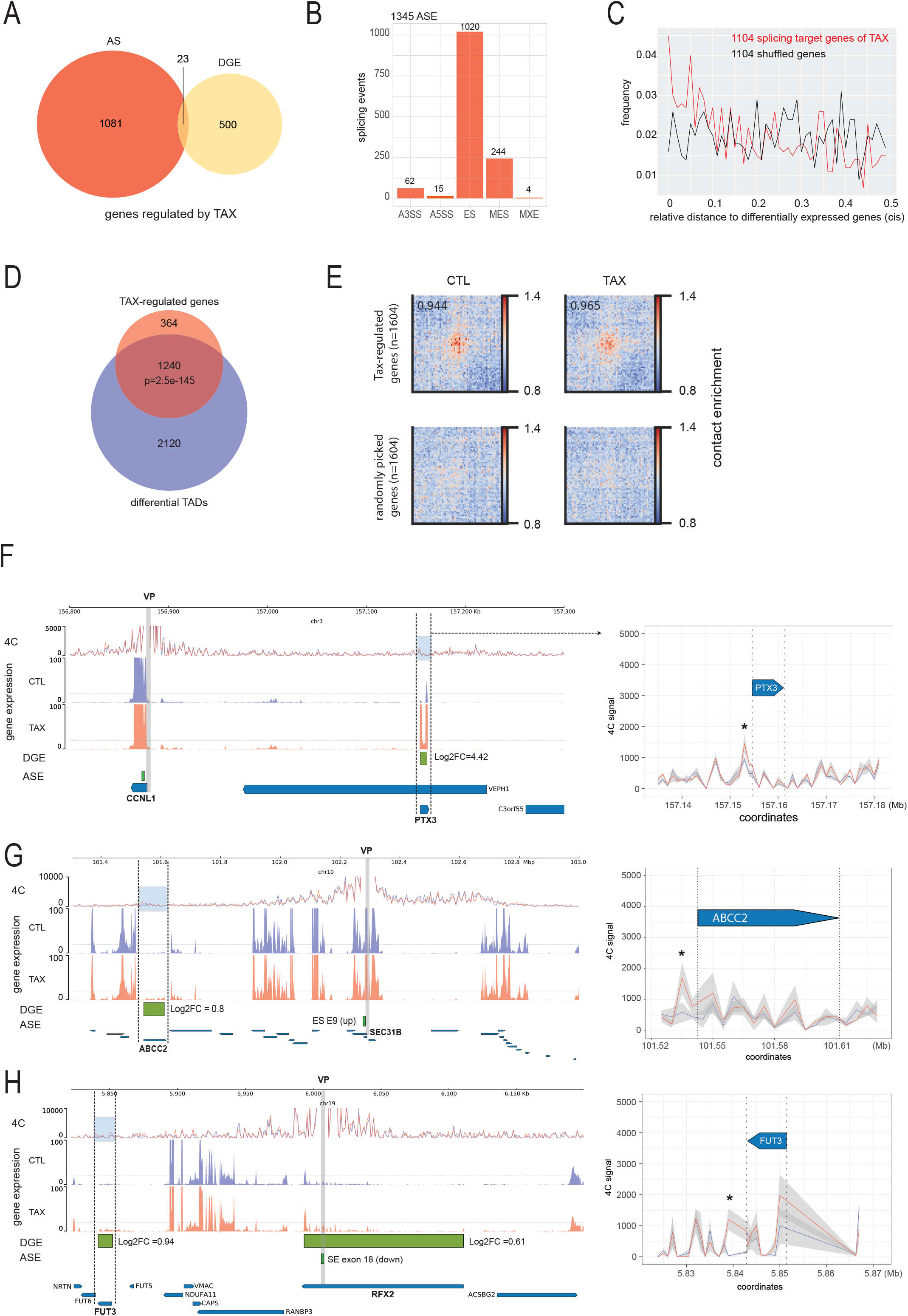
**Tax-induced 3D chromatin contacts associate with changes in both gene expression and alternative splicing.** **(A)** Venn diagram illustrating the genes affected by Tax expression in alternative splicing events (AS) and those that are differentially expressed (DGE). The gene expression threshold is set at log2fc = 0.6 with a p-value of <0.05 (glm test), and the significant threshold for □psi is set at 5% with a p-value of <0.01 (Fisher test). **(B)** Number of different types of AS events (ASE) that occur upon Tax expression. **(C)** Bedtools analysis (reldist) of the relative distance of gene submitted to alternative splicing modification to genes differentially expressed upon Tax. Control set was generated using shuffle function of bedtools that randomly permutes the genomic locations among the genome hg19. **(D)** Venn diagram displaying the number of Tax-regulated genes (DGE + ASE) that are located in differential TADs (Figure 1H). A hypergeometric statistical test was used to determine the significance of the overlap. **(E)** Pile-ups (bin size 30kb) of expected/observed signal surrounding (1Mbp) the 1604 TAX-regulated genes and 1604 randomly picked genes in CTL and TAX expressing cells. A total of 100 central pixels were used for calculating enrichment for off-diagonal pileups. (F-H) chromatin contacts evidenced in triplicated 4C analysis of *CCNL1* **(E)**, *SEC31B* **(F)**, and *RFX2* **(G)**, which were used as anchors (VP stands for viewpoint). Gene expression levels are shown for CTL (blue) and Tax(+) (red) conditions. The differentially expressed genes (DGE) and alternative splicing events (ASE) are specified above annotated genes (hg19), and Tax vs CTL values are indicated. The right panels show a zoom of the highlighted region (blue). The standard deviation across the 3 replicates of 4C assays is displayed in grey, and asterisks indicate significant changes in 4C signal (p-value <0.05, Fisher test).

**Figure 3.**
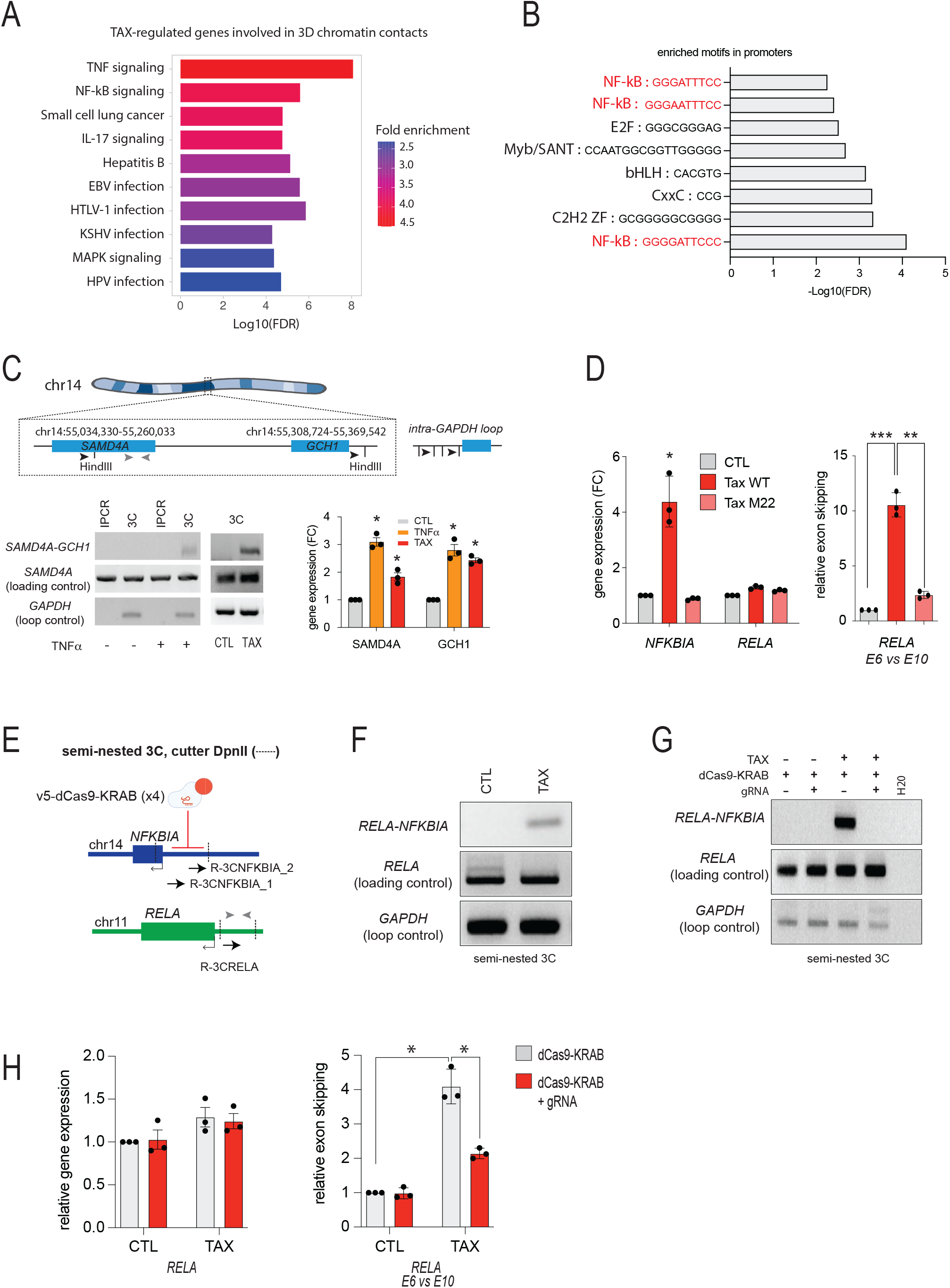
**Tax-mediated changes in the 3D genome involved contacts of NF-κB responsive genes.** **(A)** Gene Ontology (GO) analysis (KEGG pathways) of Tax-regulated genes involved in 3D chromatin contacts. The histogram indicates the significant enrichment of certain gene sets in the Tax-regulated genes (DGE+ASE). **(B)** DNA motifs enrichment analysis in the promoter regions of genes involved in 3D chromatin contacts in Tax-expressing cells using ShinyGO. **(C)** Chromosome Conformation Capture (3C) analysis of SAMD4A and GCH1 upon either 24h TNFα exposure or Tax expression is depicted. The 3C templates were prepared using HindIII, and PCR was conducted using equal weights of DNA. The resulting gels were subjected to gel electrophoresis and SYBR green staining. The presence of a band indicates a high contact frequency between the respective primer targets. Internal SAMD4A primers were used as loading control amplimers. GAPDH primers were used for amplifying an intra-GAPDH loop as previously described (34). IPCR (inverse PCR) was conducted as 3C experiments with no chromatin crosslinking steps and was used to rule out signal providing from intra-looping of HindIII-undigested fragments. The 3C products were analyzed on agarose gels stained by SYBR green. The right panel shows the relative gene expression levels induced upon TNFα exposure and Tax expression. (For all panels, asterisks correspond to statistical significance calculated using Fisher’s exact test (p-value threshold: *0.05, **0.01, ***0.001, ****0.0001). **(D)** Effects of Tax-WT and mutant Tax-M22, deficient for NF-kB activation, on gene expression levels of *NFKBIA* and *RELA* and on skipping of *RELA* exon E6. **(E)** Design of semi-nested 3C experiments targeting *NFKBIA* and *RELA*. Black arrows represent primers used for semi-nested 3C PCR, and grey arrows represent *RELA* primers used for loading control. **(F)** Semi-nested 3C analysis of chromatin contacts between *RELA* and *NFKBIA* in CTL and Tax-expressing cells. The resulting 3C products were analyzed in an agarose gel stained with Sybr-green. GAPDH primers were used for amplifying an intra-GAPDH loop as Figure 3C. **(G)** Effect of dCas9-KRAB tethered to the *NFKBIA* promoter on *NFKBIA*-*RELA* contacts. Semi-nested 3C analysis of contacts between *RELA* and *NFKBIA* was conducted in CTL and Tax-expressing cells transected with the dCas9-KRAB expression vector along with the vector expressing the four gRNAs or its empty counterpart. The resulting 3C products were analyzed in an agarose gel stained with Sybr-green. **(H)** Functional impact of dCas9-KRAB-mediated inhibition of *NFKBIA*-*RELA* contacts on gene expression levels of *RELA* andexon skipping of exon *RELA* E6.

Principal component analysis (PCA) of the Hi-C contact matrices allowed us to assess the effect of Tax on genomic compartments A and B, which correspond to euchromatin and heterochromatin regions, respectively (Figure 1B). The eigen vectors calculated from CTL and Tax samples were highly correlated (Pearson test, R=0.948, p-value <0.0001) (Sup Fig 3 C), indicating no widespread changes in genomic locations of A and B compartments between the two experimental conditions, with homotypic compartment interactions (AA and BB) being predominant (Figure 1C). However, Tax-expressing cells presented increased heterotypic compartment interactions and decreased AA contacts (Figure 1C), indicating that Tax weakens chromatin compartmentalization. Comparative analysis of compartment distribution between CTL and Tax-expressing cells revealed 13% compartment switching (A to B and B to A, Figure 1D), suggesting that the viral oncoprotein structurally affects both large heterochromatin and euchromatin regions.

Trans-chromosomal interactions have been linked to the definition of sub-compartment domains associated with distinct epigenetic regulation (50). Subcompartments A1 have been identified as highly active chromatin regions closely localized to nuclear speckles (64,65), while B1 and B2 are typically enriched in repressive marks, such as H3K27me3 and H3K9me3 (50,54). We used the CALDER algorithm (54) to infer eight subcompartments in both CTL and Tax-expressing cells (Figure 1E and Sup Fig 3 E-F). Using publicly available ChIP-seq datasets for HEK 293T cells, we confirmed that the predicted subcompartments A.1.1 and A.1.2 in control cells were markedly enriched in active marks H3K27ac and H3K4me3, while compartments B.2.1 and B.2.2 corresponded to H3K9me3 embedded regions (Sup Fig 3E). We then compared the subcompartment annotations between the two experimental conditions by considering 4 large subcompartments: A.1 (A.1.1 + A.1.2), A.2 (A.1.1 + A.1.2), B.1 (B.1.1 + B.1.2), and B.2 (B.2.1 + B.2.1). Subcompartment transitions were observed in Tax expressing cells, as demonstrated in Figure 1E and Sup Fig 3F. These transitions involved shifts from active to inactive subcompartments, with some coinciding with differential trans-chromosomal interactions (Figure 1F). Overall, these data indicated that Tax promotes a substantial reorganization of the 3D genome that likely follows dynamic chromatin regulation.

We next investigated the effects of Tax on topologically associating domains (TADs). To this end, we employed the HiC-explorer pipeline to assess the TAD-separation score, which measures the degree of separation between the left and right regions in each Hi-C matrix. We set the minimum distance between two boundaries to 50 kilobase pairs (kbp) and used publicly available CTCF binding sites for aiding identification of putative TAD boundaries (ENCFF895A0X). Our analysis revealed 3539 TADs shared by control cells and Tax-expressing cells, indicating that TAD boundaries were largely unmodified between samples (Figure 1G). Nevertheless, we observed that 40.3% and 23.4% of TADs had differential chromatin contacts at both intra- and inter-TAD levels, respectively (Figure 1H). Notably, we found a statistically significant increase in inter-TAD contacts in Tax-expressing cells (one-way ANOVA, p=0.004; Figure 1I), indicating that Tax functions as a potent modifier of host 3D chromatin architecture.

### Tax-induced 3D chromatin contacts associate with changes in both gene expression and alternative splicing

We next sought to assess the functional significance of Tax-induced 3D chromatin remodeling. Gene expression profiles including alternative splicing events of CTL and Tax(+) cells have been previously published (18). Using stringent thresholds to analyze differential gene expression (DGE) (glm-adjusted p-value < 0.05, Log2-fold change ≥ 0.6) and alternative splicing events (ASE) (p < 0.01 adjusted Fisher test, ΔPSI ≥ 0.05), we identified 523 genes with altered whole gene expression and 1,104 genes producing alternatively spliced transcripts upon Tax expression (Figure 2A). A total of 1345 Tax-induced alternative splicing events was identified, consisting largely in skipped exon (SE) (Figure 2B). In line with previously published data (18,23), a minority of Tax-regulated genes (5%, 23/523) was affected at both gene expression and splicing levels (Figure 2A), suggesting that Tax-induced transcription and alternative splicing are two distinct and complementary mechanisms that profoundly modify the host transcriptome upon HTLV-1 infection. We examined the relative 2D/linear distribution of Tax-regulated genes across the genome and found that genes producing alternatively spliced transcripts tended to be located closer to differentially expressed genes than expected by chance, suggesting that, while p65/RelA-driven alternative splicing and promoter activation of individual genes may operate independently (18), these two mechanisms could be locally and spatially interrelated (Figure 2C). The coordinates of Tax-regulated genes (DGE + ASE) significantly overlapped with TADs that exhibit differential contacts (Figure 2D), leading us to examine whether Tax-induced gene regulations could colocalize with differential 3D chromatin interactions. To this end, we cross-compared the differential Hi-C contact matrix with genome coordinates of Tax-regulated genes and found that the majority of Tax-regulated genes appear to be intertwined in a highly interconnected network of 3D chromatin contacts that occur at both cis-chromosomal and trans-chromosomal levels (Sup Fig 4). Using pile-up analysis to quantify the overall interaction strength between genomic loci of TAX-regulated genes, we confirmed their specific enrichment in contacts in both CTL and TAX expressing cells, while no such enrichment was observed with randomly selected genes (Figure 2E). Cells expressing TAX showed a modest increase in the score of contact enrichment, suggesting that TAX may strengthen some interactions among its regulated genes. In line with this, using differential Hi-C matrix binned to 30 kbp, we identified 73% deregulated genes (including both DGE and ASE, 1173/1604) distributed in 1451 differential 3D chromatin interactions (Log2FC ≥ 0.5) (Sup Fig 4). Of those, 237 chromatin contacts involved genes affected in expression, consistent with previously described transcriptional condensates clustering multiple genes co-regulated in transcription (34,35,39). Additionally, 1047 chromatin contacts involved genes undergoing alternative splicing, of which 731 were spatially gathered with genes differentially expressed, suggesting that multigene complexes might coordinate both transcription and alternative splicing of co-regulated genes (Sup Fig 4).

**Figure 4.**
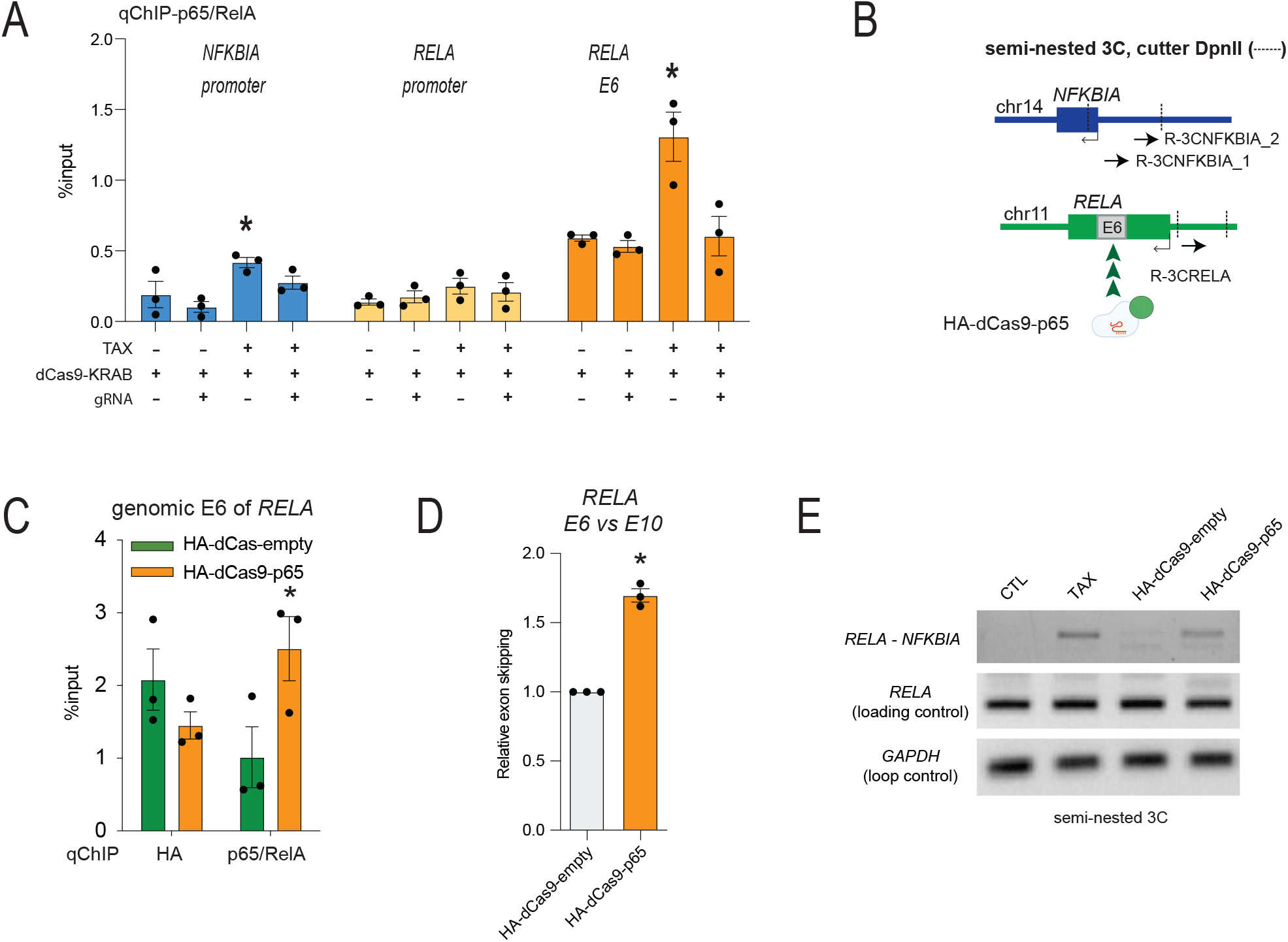
**Enhanced exon skipping of *RELA* E6 and increased chromatin contacts between *RELA* and *NFKBIA* upon p65/RelA recruitment to the gene body of *RELA*.** **(A)** Quantitative ChIP analysis of p65/RelA occupancy of *NFKBIA* promoter, *RELA* promoter, and genomic exon E6 of *RELA* in cells with and without Tax expression, and with or without disruption of *NFKBIA*-*RELA* contacts by dCas9-KRAB. Histograms show the mean fold enrichment relative to input DNA, with error bars indicating SEM. **(B)** Design of semi-nested 3C experiment for analyzing *NFKBIA*-*RELA* contacts in cells expressing dCas9-p65/RelA tethered to the genomic exon E6 of *RELA* using 4 distinct gRNAs. **(C)** Validation of dCas9-p65/RelA tethering to the genomic exon *RELA* E6 by qChIP against the HA tag and p65/RelA. HEK cells were transfected with the expression vector of HA-dCas9-p65/RelA or the empty vector expressing HA-dCas9-empty (with no p65/RelA fused). **(I)** Effect of genomic *RELA* E6-tethered dCas9-empty and dCas9-p65/RelA on exon skipping of *RELA* E6. RT-qPCR analysis was performed using RNA extracted from cells expressing HA-dCas9-empty or HA-dCas9-p65/RelA and with or without Tax expression. **(D)** Effect of tethered dCas9-empty and dCas9-p65/RelA into genomic exon E6 on gene-gene contacts between *NFKBIA* and *RELA*. Semi-nested 3C experiments were carried out in cells expressing dCas9-p65/RelA tethered to the genomic exon E6, as illustrated in panel B. Cells with and without Tax expression were used as controls. For all panels, asterisk correspond to statistical significance assessed using the Fisher test (*p < 0.05).

To further improve this Hi-C prediction, we conducted 4C assays to investigate physical interactions between genes expressing alternative spliced isoforms and distant genes that were concomitantly deregulated at the gene expression level by Tax. We selected three distinct viewpoints in genes *CCNL1*, *SEC31B*, and *RFX2*, which were identified above as Tax-regulated targets for alternative splicing of exons E4, E9, and E18, respectively (Figure 2F-H) (18). Using these viewpoints in triplicate 4C assays, we were able to ensure that the chromatin contact frequencies remained reliable and consistent across different replicates within each experimental condition (as illustrated in right panels (grey lines) of Figures 2F-H). The comparison of the contact landscapes of each viewpoint between cells expressing Tax and control cells indicated that the genes *CCNL1*, *SEC31B*, and *RFX2* exhibited a higher frequency of interactions with distal genes that were upregulated by Tax. More specifically, *CCNL1* showed increased connections with *PTX3* located 295 kbp downstream, *SEC31B* with *ABCC2* at a distance of 750 kbp, and *RFX2* with *FUT3* positioned 268 kbp downstream (Figure 2F-H). Overall, these results provide compelling evidence that Tax triggers long-range chromatin interactions that involve genes undergoing differential expression and/or alternative splicing regulation.

### Tax-induced NF-κB activation promotes gene-gene contacts

Because previous work had highlighted a central role of the NF-κB pathway in Tax-regulated transcription and alternative splicing, we hypothesized that this pathway could also promote the 3D chromatin interactions described above. We thus assessed whether specific gene subsets, including NF-κB-regulated genes, were enriched in chromatin interactions generated by Tax. Gene ontology analysis of Tax-regulated genes (including both DGE and ASE) identified in 3D chromatin contacts indeed pointed to a significant enrichment in TNF-α and NF-κB signaling pathways, while remaining terms were related to oncoviral infection including that of HTLV-1 (Figure 3A). In line with this, analysis of transcription factor motifs in promoter sequences involved in chromatin contacts revealed a significant enrichment in NF-κB motifs (Figure 3B), suggesting a potential link between NF-κB signaling, alterations in 3D chromatin structure, and gene regulation in response to TAX. These results were reminiscent of previous studies reporting the formation of multigene complexes in which interacting genes are especially activated by NF-κB upon TNF-α stimulation (34,35). Accordingly, we sought to assess by chromosome conformation capture (3C) whether such NF-κB multigene complexes also occurred in Tax expressing cells. To this end, we developed a 3C assay and an inverse PCR (IPCR) assay (with no crosslinked chromatin) used as negative control. Validating our approach and ruling out false positive products derived from ligation of free DNA fragments, the 3C analysis, but not IPCR assay, detected a previously described TNF-α-related gene-gene interaction between *SAMD4A* and *GCH1* genes (∼0.3 Mbp distance on chromosome 14) (Figure 3C) (34). Similar results were obtained in cells expressing Tax, indicating that Tax mimics TNF-α-activated cis-contacts (Figure 3C). In both experimental conditions, the *SAMD4A*-*GCH1* interaction coincided with a significant increase in *SAMD4A* expression (Figure3C, right panel), thereby suggesting that, as described for TNF-α, Tax promotes the formation of multigene complexes in which genes are coregulated in transcription.

Building on previous evidence of trans-chromosomal contacts between chromosomes 11 and 14 induced by TNF-α (34,35), we further investigated whether these contacts might also involve genes regulated by Tax. Given their pivotal roles in NF-κB signaling, viral persistence, and ATL development,we focused our attention on *NFKBIA* (chr14:35870716-35873960) and *RELA* (chr11:65421067-65430443) that encode the inhibitor IκBα and the effector p65/RelA of NF-κB signaling, respectively (66). Of note, among the numerous genes encoding members of the TAX protein-protein interactome and often affected by ATLL-related mutations (65,66), only *NFKBIA* and *RELA* were found in chromosomes 11 and 14 (66,67). RNA-seq data showed that *NFKBIA* was significantly upregulated upon Tax expression, while *RELA* displayed no significant changes in gene expression but was regulated at the alternative splicing level by the exon skipping of exon 6 (Sup Table 3). We confirmed these RNA-seq predictions by RT-qPCR assays in HEK cells expressing Tax as well as in HTLV-1-transformed C91PL cells (Figure 3D and Sup Fig 5A). Of note, the protein level of p65/RelA in HEK cells appeared unaffected by Tax, whereas that of IκBα was downregulated, which was consistent with its proteasomal degradation described upon Tax expression (Sup Fig 5B) (68,69). In contrast to the wild-type Tax, the ectopic expression of TaxM22, a mutant deficient for NF-κB activation, failed to influence both *NFKBIA* expression and *RELA* exon skipping (Figure 3D), indicating that both transcriptomic events affecting *NFKBIA* and *RELA* upon Tax expression relied on NF-κB activation.

We next sought to test whether the concomitant deregulation of *NFKBIA* and *RELA* corresponded to increased trans-chromosomal contacts between chromosomes 11 and 14 upon Tax expression. To this end, we developed semi-nested 3C assays using both promoters of *RELA* and *NFKBIA* as anchors (Figure 3E). This approach revealed a single PCR signal attributed to *RELA*-*NFKBIA* contacts in Tax expressing cells, but not in CTL cells (Figure 3F). In comparison, there was no change in PCR signals deriving from an intragenic fragment of *RELA* used as loading control, as well as from an intra-GAPDH chromatin loop used as 3C control (34). These data demonstrated that Tax induced chromatin contacts between *NFKBIA* and *RELA* genes localized on chromosomes 14 and 11, respectively.

To determine whether the formation of a Tax-induced multigene complex involving *NFKBIA* and *RELA* participates in regulating the alternative splicing of *RELA* transcripts, we aimed to examine the *RELA* E6 skipping event in cells that are impaired for *NFKBIA*-*RELA* interaction. To this end, we used the dCas9-KRAB system (70), already identified as an effective steric barrier of chromatin contacts (71), to disrupt the contact between *NFKBIA* and *RELA* genes. Four distinct guide RNAs were designed to tether the dCas9-KRAB construct to the *RELA*-interacting genomic region of *NFKBIA* (Figure 3E). Quantitative ChIP (qChIP) experiments confirmed the specific tethering of dCas9-KRAB onto the *NFKBIA* target when compared to a gRNA-free experiment (Sup Figure 5C). Semi-nested 3C assays demonstrated that the influence of Tax on the *RELA*-*NFKBIA* interaction was compromised under these conditions (Figure 3G). Tethering the dCas9-KRAB on *NFKBIA* promoter did not interfere with the expression level of *RELA* but it significantly reduced the effect of Tax on *RELA* E6 exon skipping (Figure 3H). Importantly, we ascertained that the overexpression of ectopic IκBα had no impact on this negative effect of dCas9-KRAB (Sup Fig 5D-E), indicating that the loss of Tax-induced skipping of *RELA* E6 through dCas9-KRAB tethering to *NFKBIA* is due to disrupted contacts between *NFKBIA* and *RELA* rather than a reduced cellular abundance of IκBα. Collectively, these data define trans-chromosomal contacts between chromosomes 11 and 14 as an underlying mechanism for the concomitant Tax-driven regulation of *NFKBIA* expression and alternative splicing of *RELA* transcripts.

### Intragenic p65/RELA occupancy of *RELA* leads to *RELA*-*NFKBIA* contacts and alternative splicing of *RELA* E6

In light of our previous findings that p65/RelA occupancy of genomic exons can influence gene expression through alternative splicing (18), we next conducted qChIP assays to assess the chromatin distribution of p65/RelA across *RELA* and *NFKBIA* in cells with or without Tax expression. As shown in Figure 4A, Tax induced an average 2-fold enrichment of p65/RelA binding on the *NFKBIA* promoter, whereas it had no significant effects on *RELA* promoter occupancy. These data reflected the effects of Tax on the gene expression levels of *NFKBIA* and *RELA* (Figure 3D). In addition, we observed a significant increase in p65/RelA occupancy at the genomic exon E6 of *RELA* of Tax-expressing cells compared to control cells, suggesting that exon E6 belongs to exon subsets that are regulated by p65/RELA (Figure 4A). Moreover, disrupting *NFKBIA*-*RELA* contacts with the dCas9-KRAB in Tax-expressing cells led to a marked reduction in p65/RelA recruitment at the *NFKBIA* promoter, along with a significant decrease in p65/RelA occupancy of genomic exon E6, while there was no significant impact on p65/RelA occupancy of the *RELA* promoter (Figure 4A). These findings suggest a direct relationship between p65/RelA, alternative splicing, and gene-gene contact regulation.

We previously demonstrated that experimental chromatin enrichment of p65/RelA in proximity to its genomic exon target can effectively lead to alternative splicing modification (18). To further investigate the role of p65/RelA in alternative splicing and 3D chromatin contacts, we designed dCas9 constructs fused or not to p65/RelA to assess the effect of p65/RelA locally tethered to the genomic *RELA* E6 (Figure 4B). We validated the specific tethering of the dCas9 constructs to this region using qChIP assays, compared to a dCas9-empty construct (Figure 4C). At the RNA level, this dCas9-mediated local p65/RelA enrichment resulted in exon skipping of E6 (Figure 4D), confirming that *RELA* E6 is an NF-κB responsive exon. Additionally, semi-nested 3C analysis of *NFKBIA*-*RELA* interactions revealed that E6-tethered dCas9-p65/RelA was sufficient to promote gene-gene interactions between *NFKBIA* and *RELA* (Figure 4E), thus identifying p65/RelA as an effector of multigene complex formation where NF-κB responsive genes are coordinated in their regulation at both transcriptional and alternative splicing levels.

## DISCUSSION

The 3D genome conformation is crucial for gene regulation, and understanding how viruses manipulate it can reveal fundamental principles of gene regulation and cell behavior. Here, we present evidence that the HTLV-1 viral oncoprotein Tax remodels the 3D chromatin architecture of its host cells. Using Hi-C and 4C analyses, we found that Tax-induced changes in long range chromatin interaction frequencies, compartmentalization, and inter-TAD interactions coincided with differential gene expression and alternative splicing. We demonstrated that gene-gene interactions led to coordinated splicing and transcriptional events upon NF-κB activation. Our data show that Tax-induced NF-κB activation promotes contacts between *NFKBIA* and *RELA* genes, which coincides with the skipping of exon E6 of *RELA* and upregulation of *NFKBIA* expression. Disrupting the *NFKBIA*-*RELA* interaction impacted *RELA* E6 exclusion at the RNA level, while targeted chromatin recruitment of dCas9-p65/RelA triggered *NFKBIA*-*RELA* interaction and *RELA* E6 skipping. Our findings suggest that p65/RelA plays a role in assembling multigene complexes, coordinating transcription and alternative splicing regulation of NF-κB-responsive genes. This highlights a new mechanism coordinating transcriptional and post-transcriptional changes through the NF-κB pathway and shows how HTLV-1 Tax impacts molecular mechanisms linking 3D genome conformation to transcriptome diversity.

Numerous cellular signaling pathways are known to influence alternative splicing (72,73), in addition to their effects on gene transcription. However, the specific mechanisms responsible for fine-tuning cellular responses to stimuli are not yet fully understood. In this context, studies examining the dynamics of 3D genome conformation have revealed the importance of multigene complex formation in gene coregulation upon cell signaling (34,35,40), where long-range interactions between cotranscribed genes help to organize transcription within nuclear condensates by establishing hierarchical relationships between gene loops. In light of our findings, we propose a model in which Tax-induced NF-κB activation leads to a persistent accumulation of p65/RelA at promoters and enhancers, forming nuclear condensates that assemble NF-κB-responsive genes for coordinated transcription, in line with previous reports (34,35,40). Simultaneously, based on the same physicochemical properties, genes with p65/RelA bound to genomic exons might be attracted to NF-κB factories without any consequence on their transcriptional activity, due to the absence of NF-κB factors at their promoters. However, these multigene complexes could create a suitable environment for facilitating p65/RelA-dependent alternative splicing regulation.

Accordingly, the exon target specificity and the resulting splicing outcome may then depend on the p65/RelA protein-protein interactome, in which the presence of various RNA helicases like DDX17, DDX5, DHX9, DDX3X, G3BP1, DHX15, DDX1, and DDX21 is notable, as well as other splicing regulators including SF3B130 (19) and FUS (22,74). Interestingly, these factors have been recently linked to regulatory splicing subnetworks associated with genes located in the center of nucleus and composed of GC-rich intron-exon units (45). This observation is consistent with the features of splicing events initiated by Tax, which predominantly affect GC-rich exons (18). As an additional layer, splicing-associated chromatin signatures (SACS) have been shown to potentially recruit distinct splicing regulators to pre-mRNA of genes within the same functional pathway (75,76). Given its capacity to recruit diverse chromatin remodelers and splicing factors, p65/RelA emerges as a strong candidate for facilitating SACS. Since histone modifications influence both local and global 3D genome organization, p65/RelA could act as a key initiator of intricate and interconnected mechanisms that govern gene movements and splicing decisions upon NF-κB activation. Future studies should examine these points, focusing on characterizing the p65/RelA interactome and its role in alternative splicing regulation in the context of 3D genome.

Although the long-term effects of Tax-induced NF-κB activation have been widely studied, it remains unclear whether this activation leads to unique expression patterns of NF-κB-responsive genes that are specific to Tax. Our study reveals that Tax-induced NF-κB activation leads to gene-gene contact-dependent regulation of alternative splicing for exon E6 of *RELA* in both HEK and HTLV-1 transformed cells, resulting in the expression of a spliced isoform lacking E6. This isoform encodes a shorter p65/RelA protein of approximately 44 kDa (p65/RelAΔE6), which, unlike its wild-type counterpart, does not independently alter the expression levels of NF-κB-responsive genes. However, when Tax is expressed, ectopic expression of p65/RelAΔE6 appears to repress A20 and BCL3 while upregulating IL8 (Sup Fig 5F-H). These findings suggest that *NFKBIA*-*RELA* contacts upon Tax-induced NF-κB activation allow the generation of a *RELA* spliced isoform that might contribute to refining Tax activation-specific gene expression profiles.

Recent studies have reported that other oncogenic viruses, including human papillomavirus (HPV) (77), Epstein-Barr virus (EBV) (78), and hepatitis B virus (HBV)(79), reshape the 3D structure of their host cell genome, potentially contributing to complex mechanisms underlying gene perturbations that lead to cancer development. Regarding HTLV-1, its integration has been demonstrated to alter the 3D genome structure by forming loops between the proviral CTCF binding site and host genome regions located in cis (80). It has been proposed that the integration of an HTLV-1 provirus into a genomic region that typically occupies a transcriptionally repressive compartment in the nucleus, such as lamina-associated domains (LADs), could favor the survival of HTLV-1-infected cellular clones in vivo (81). However, some gene promoters associated with the nuclear lamina have been shown to escape transcription repression (82), suggesting a more complex and dynamic regulation in peripheral region of the nucleus. Given our findings that Tax can promote changes in long-range chromatin interactions affecting both cis- and trans-chromosomal regions, future investigations should examine how bursts of Tax expression influence the magnitude of proviral-genome contacts and their consequences on the clonal selection of infected cellular clones and the development of HTLV-1-associated diseases.

In summary, our study demonstrates that the HTLV-1 viral oncoprotein Tax remodels the 3D chromatin architecture of its host cell, influencing gene-gene interactions and coordinating transcription and alternative splicing events upon NF-κB activation. This sheds new light on the orchestration of the interplay between 3D genome conformation and transcriptome diversity, building upon earlier works from our group and others that demonstrated the role of intragenic chromatin loops in regulating transcription and alternative splicing (41,83–85). Our findings raise the question whether similar mechanisms could pertain to other cellular signaling pathways, such as TGFβ that has been linked to TGFβ factories and splicing regulation through a SMAD3:HNRNPE1 axis (34,86). Further investigations are needed to fully understand the complex relationship between cellular signaling pathways, chromatin architecture, coordinated transcriptional and alternative splicing regulation, and the impact of HTLV-1.

## DATA AVAILABILITY

The raw data for 4C, Hi-C, and RNA-seq have been deposited in NCBI’s Gene Expression Omnibus and are accessible through GEO Series accession number GSE233887, GSE233888, and GSE123752, respectively.

## ACKNOWLEDGEMENTS

We gratefully acknowledge support from Laurent Modolo at the LBMC for bio-informatic advice, and from the PSMN (Pôle Scientifique de Modélisation Numérique) of the ENS de Lyon for computing resources. FM thanks Chloé Journo, Daniel Jost, and Aurele Piazza for their valuable discussions and insightful comments.

## FUNDING

This work was supported by the Ligue Contre le Cancer (Comité de du Rhône), the Fondation ARC (ARC PJA20151203399), and the Agence Nationale pour la Recherche (program INFLASPLICE). P.M. was supported by a bursary from the French Ministry of Higher Education and Science, the Fondation de France, and ANR INFLASPLICE; J.L., M.B., and L.B.A. by the Ligue Contre le Cancer; N.F by ENS-Lyon; and T.S., C.F.B, D.A., and F.M. by INSERM.

### CONFLICT OF INTEREST

The authors declare no competing interests.

**Supplementary Tables**

**Supplementary Table 1: list of oligonucleotides**

**Supplementary Table 2: comprehensive Overview of Interaction Data**

**Supplementary Table 3: Whole gene expression and alternative splicing changes upon Tax expression.** Whole gene expression threshold was set to log2(FC)=0.6, p<0.05 (glm and Wald test). For differential splicing, deltaPsi threshold was set to 0.5 p<0.01 (Fisher test). These data have been already published in Ameur LB, *et al.* in Nature Communications 2020 (18).

## SUPPLEMENTARY DATA

Supplementary Data are available at NAR online.

**Supplementary Figure 1.**
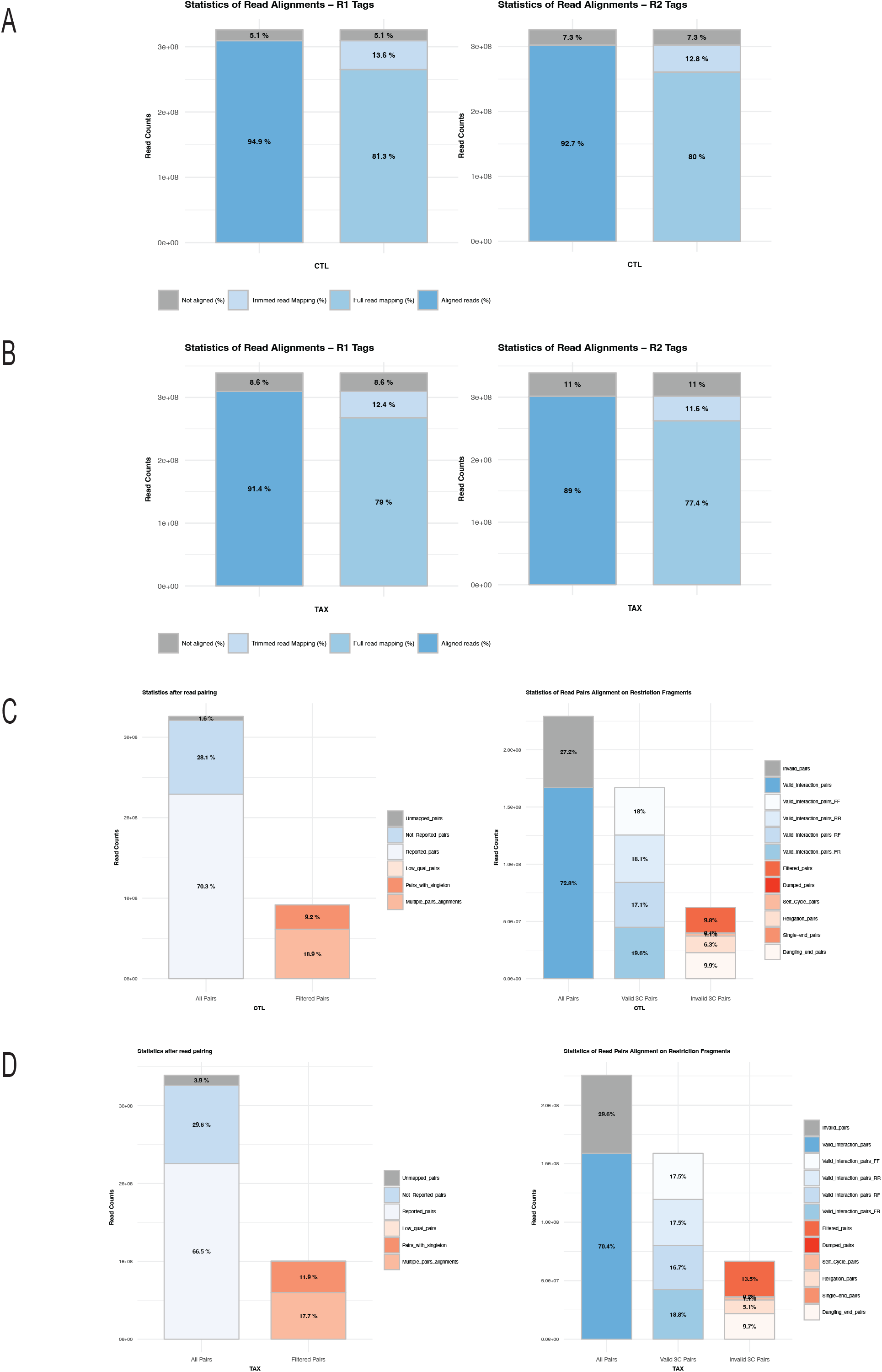
Hi-C data quality control analysis by HiC-Pro pipeline for Tax(+) and CTL conditions. (A-B) Quality control of read alignment and pairing for the CTL and Tax conditions, respectively. During this step, low-quality alignment, singleton, and multiple hits are removed. (C-D) represent the read pair filtering step, where read pairs are assigned to a restriction fragment and invalid pairs such as dangling-ends and self-circles are tracked but excluded from downstream analysis.

**Supplementary Figure 2.**
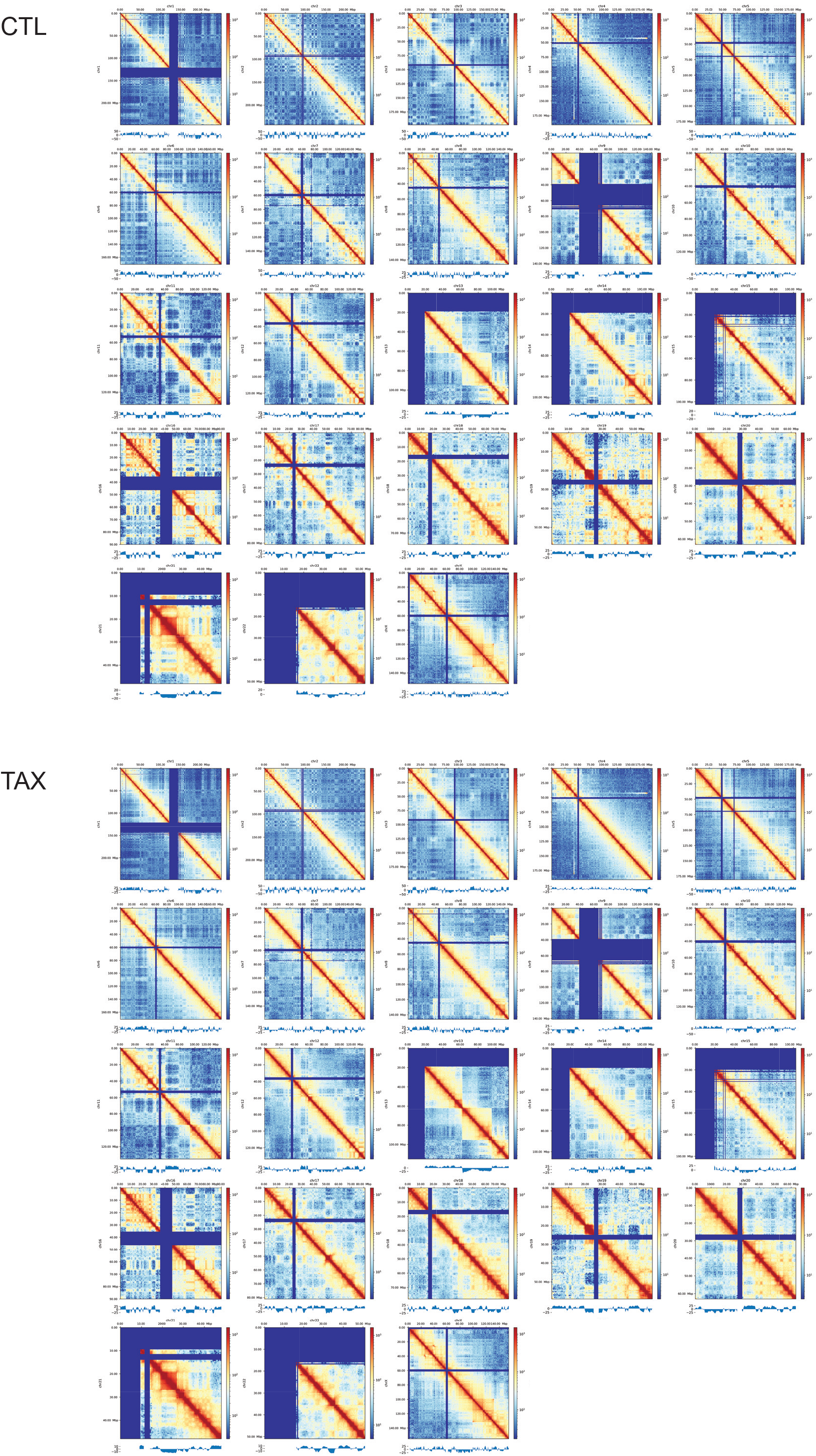
Hi-C heatmaps and compartment analysis at 500kbp resolution. The figure shows Hi-C contact maps for each chromosome, visualized in logarithmic scale. Each heatmap represents the contact frequencies between genomic loci at a resolution of 500kbp. Below each heatmap, the eigen vector value of the corresponding chromosomal segment is presented. This value characterizes the compartmentalization of the genome into A (positive) and B (negative) compartments, which correspond to open and closed chromatin, respectively. The data were preprocessed and normalized using ICE algorithm. The visualization was generated using hicexplorer software.

**Supplementary Figure 3.**
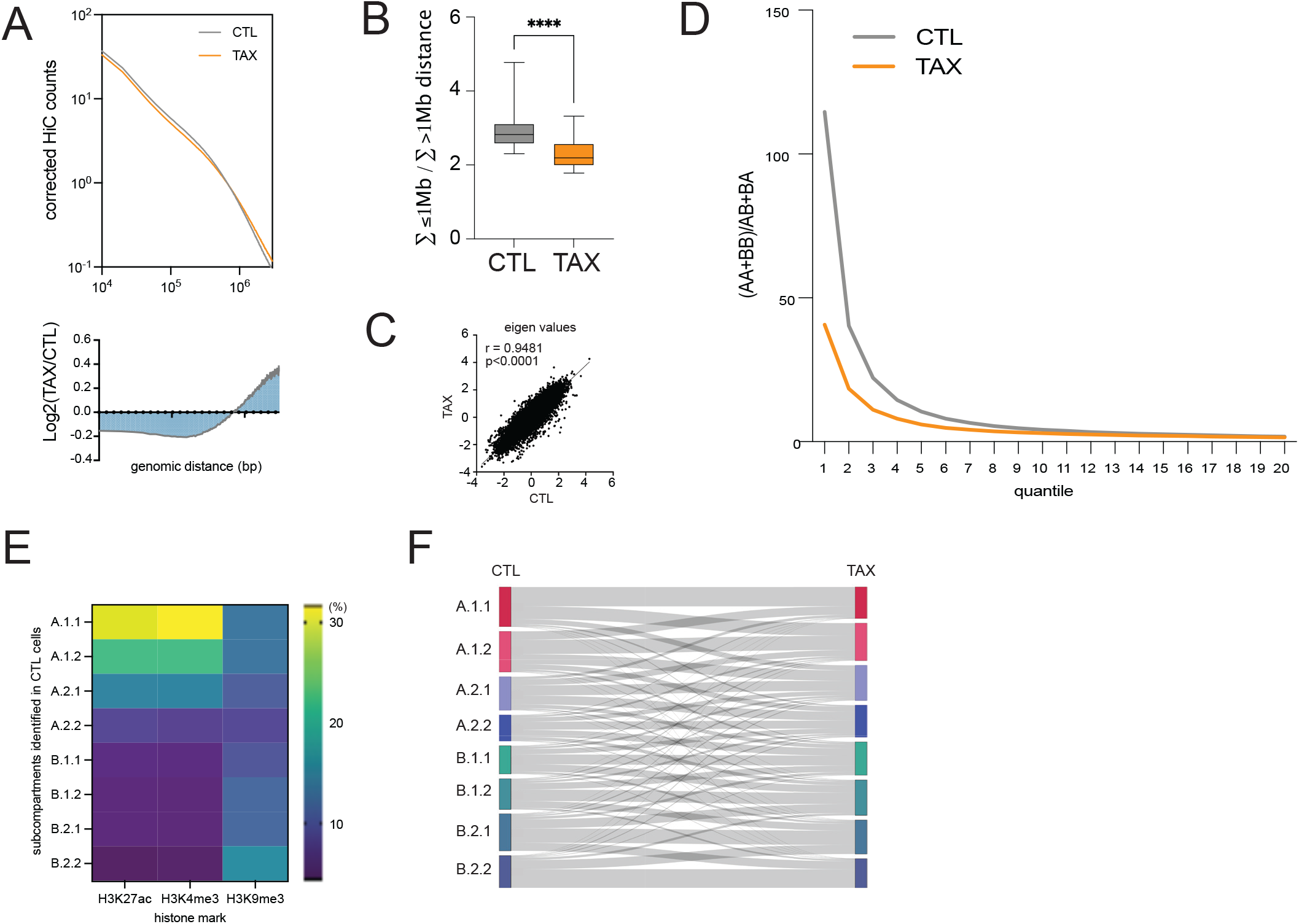
Tax alters long range chromosomal interaction and the compartmentalization of the genome. **(A)** Contact probability was calculated using hicexplorer as a function of genomic distance for interactions within individual chromosome arms across the whole genome. Lower panel indicates the log2FC of differential contact frequency between Tax and CTL condition. **(B)** The ratio of short-range (≤1 Mb) versus long-range (≥1 Mb) interactions **(SVL)** was calculated for each chromosome arm across the whole genome using hicexplorer. The SVL is significantly lower in Tax-expressing cells than in CTL cells (****P < 0.0001). **(C)** cis-eigen values are not affected genome wide by the expression of Tax Pearson test, R=0.948, p-value <0.0001). **(D)** Compartmentalization strength in CTL and Tax expressing cells. The global strength for compartmentalization is computed genome-wide as (AA + BB) / (AB + BA) after rearranging the bins of obs/exp based on their corresponding eigen values. **(E)** Correlation of 8 subcompartments identified by CALDER in CTL cells and histone mark enrichment for H3K27ac, H3K4me3, H3K9me3 obtained from publicly available Chip-seq data in HEK cells. **(F)** Sankey plot showing subcompartments switching upon Tax expression.

**Supplementary Figure 4.**
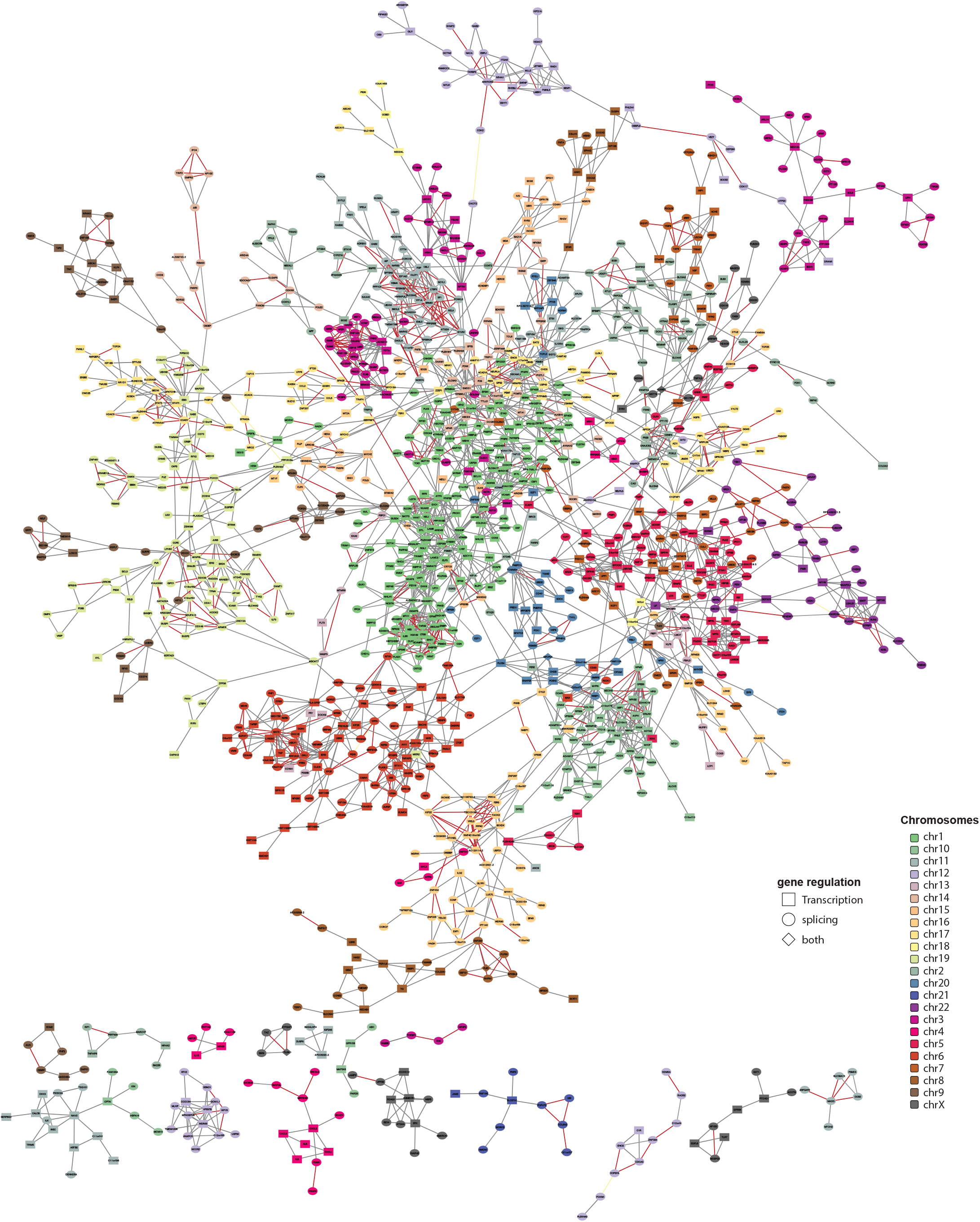
Network analysis of Tax-regulated genes (DGE and ASE) involved in 3D chromatin contacts. Overall, 1369 genes were identified to be involved in 3D chromatin contacts in Tax expressing cells, forming a main network consisting of 1247 Tax-regulated genes that are connected through 2671 chromatin contacts. Genes that belong to the same chromosome are color-coded uniformly. The shape of nodes representing genes indicates their modification upon Tax expression. Red edges indicate differential contact frequency between Tax expressing cells and CTL cells (abs(Log2FC)>0.5).

**Supplementary Figure 5.**
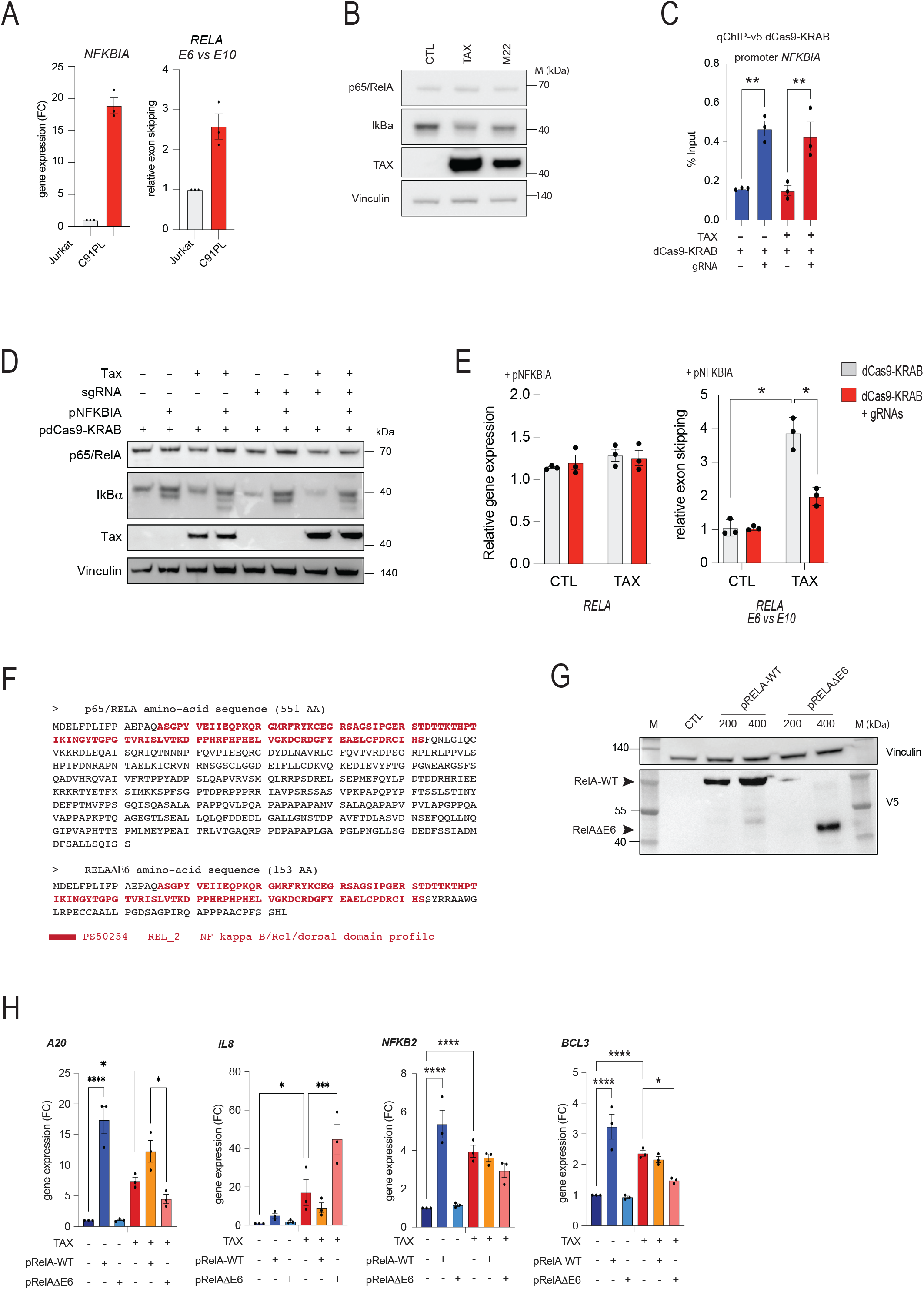
DNA binding of p65/RelA to the genomic exon E6 of *RELA* leads to concomitant *RELA*-*NFKBIA* interactions and exon skipping of *RELA* E6. **(A)** Quantitative RT-PCR analysis of gene expression levels of **NFKBIA** and **RELA**, as well as E6 exon skipping of **RELA** in Jurkat cells and HTLV-1 transformed T-cells C91PL. **(B)** Western blot analysis of p65/RelA, IκBα, and Tax in cells expressing Tax, Tax M22, and control cells. Vinculin was used as a loading control. **(C)** Validation by qChip of dCas9-KRAB tethering using four gRNAs (see Methods) on the **NFKBIA** promoter. The percentage input is presented for cells expressing Tax and those not expressing Tax. **(D)** Western blot analysis of p65/RelA, IκBα, and Tax in control cells and Tax-expressing cells transfected with pdCas9-KRAB along with the expression or not of guide RNAs and the vector pNFKBIA encoding IκBα. Vinculin was used as a loading control. **(E)** The ectopic expression of IκBα does not affect the effect of Tax, and that of **RELA** E6-tethered dCas9-KRAB, on exon skipping of **RELA** E6 (compared to Figure 4D). Quantitative RT-PCR analysis was carried out as described in Figure 3C. **(F)** Prediction of the protein encoded by the splicing isoform of **RELA** skipped for exon E6. The GenBank sequence NM_021975 was used as reference for the full transcript of **RELA**. An open-reading frame prediction using Expasy tools revealed that, in comparison to the wild-type isoform that encodes 551 amino acids, the mRNA isoform lacking exon E6 - referred to as RelAΔE6 - encodes a truncated protein isoform of p65/RelA that consists of 153 amino acids (predicted at 44 kDa). This truncated isoform shares the rel homology domain with its wild-type counterpart but lacks other domains that are involved in DNA binding and protein-protein interactions. **(G)** Western blot analysis of of RelA-WT and RelAΔE6 constructs using V5 antibodies. HEK293T cells transiently transfected with 200ng and 400ng of expression vectors for p65/RelA-WT (pRelA-WT) and p65/RelAΔE6 (pRelAΔE6) or empty vectors. **(H)** Effect of ectopic expression of p65/RelA-WT and the splicing isoform p65/RelAΔE6 on gene expression levels of A20, IL8, NFKB2 and BCL3. HEK cells were transiently transfected with 400 ng of expression vectors for p65/RelA-WT (pRelA-WT) and p65/ (pRelAΔE6) (or empty vector as control) along with Tax expression vector or its empty vector control. Total RNA was extracted from the cells at 48 hours post-transfection, and RTqPCR was performed as described in the Methods section.

